# TBtools - an integrative toolkit developed for interactive analyses of big biological data

**DOI:** 10.1101/289660

**Authors:** Chengjie Chen, Hao Chen, Yi Zhang, Hannah R. Thomas, Margaret H. Frank, Yehua He, Rui Xia

## Abstract

The rapid development of high-throughput sequencing (HTS) techniques has led biology into the big-data era. Data analyses using various bioinformatics tools rely on programming and command-line environments, which are challenging and time-consuming for most wet-lab biologists. Here, we present TBtools (a <u>T</u>oolkit for <u>B</u>iologists integrating various biological data handling <u>tools</u>), a stand-alone software with a user-friendly interface. The toolkit incorporates over 100 functions, which are designed to meet the increasing demand for big-data analyses, ranging from bulk sequence processing to interactive data visualization. A wide variety of graphs can be prepared in TBtools, with a new plotting engine (“JIGplot”) developed to maximum their interactive ability, which allows quick point-and-click modification to almost every graphic feature.

TBtools is a platform-independent software that can be run under all operating systems with Java Runtime Environment 1.6 or newer. It is freely available to non-commercial users at https://github.com/CJ-Chen/TBtools/releases.

## The Rationale for TBtools development

### Developed for wet-lab biologists

The exponential growth of biological data has come with the rapid development and renovation of high-throughput sequencing (HTS) techniques. Managing big-data and decoding the underlying bio-information effectively and efficiently presents a significant challenge to wet-lab biologists. Various bioinformatics software, pipelines, and packages have been developed to meet this challenge, however, most of these tools are packaged as scripts written in disparate programming languages, and require a working knowledge of the command-line environment. This lack of easily accessible tools remains a significant obstacle for wet-lab biologists who want to process their own data but lack proficient computational skills. HTS technologies are frequently used to investigate biological phenomena on a genomic scale. Unfortunately, the big-data generated is often underutilized when experimental biologists run into programming roadblocks. Here, we present TBtools, a <u>T</u>oolkit for <u>B</u>iologists integrating various biological data handling <u>tools</u> with a user-friendly interface; out aim is to accelerate discoveries by providing an out-of-the-box solution to the data-handling dilemma of biologists. TBtools contains an extensive collection of functions, which integrate into a graphic user interface (GUI) that can be easily navigated using point-and-click icons. For each function in TBtools, we designed its GUI panel according to the most straightforward IOS logic, i.e., Set <u>I</u>nput Data, Set <u>O</u>utput Path if Required and Click <u>S</u>tart Button. This interface makes the handling of big-data a more pleasant and efficient experience.

### Developed as an integrative toolkit

In order to handle large biological data, researchers are currently required to work under a command-line environment, and use several, even dozens of independent tools sequentially. For instance, to identify homologous genes of a specific gene family from a species, users must access genomic data which are commonly available as two separate files, a genome sequence file in FASTA format and a gene structure annotation file in GFF3/GTF format; then they should firstly invoke “gffread” (Trapnell et al., 2013) which only works under Unix-like operating system to retain gene sequences, secondly apply BLAST program (Camacho et al., 2009) to get an ID list of homologous genes, and finally use home-brew scripts or other tools like “seqkit” (Shen et al., 2016) to extract sequences of homologous genes. In TBtools, we aim to integrate all of these functions in a Java Run Time Environment (version >= 1.6), which is compatible across the three major operating systems (Windows, Machintosh, Unix). All of the functions can be implemented through simple point-and-click using a mouse. Sequential steps for data analyses and visualizations have been integrated into a single IOS workflow. Functions in TBtools are, in most cases, coded from scratch using Java or achieved by invoking cross-platform programs (e.g., BLAST). To date, more than 100 functions are available in TBtools, covering the most commonly used tools for bioinformatic analyses. These include big-data preview, data format conversion, basic sequence management, interactive data visualization in numerous forms that span from simple Venn diagrams to sophisticated synteny plots (Fig. 1). Notably, the development of TBtools was highly collaborative and greatly motivated by the true needs of wet-lab biologists. In the past five years, the tool has attracted over 15,000 stable users. Many of these users actively provide informative feedback and suggestions, which has significantly enhanced the functionality and features of TBtools.

**Figure 1.**
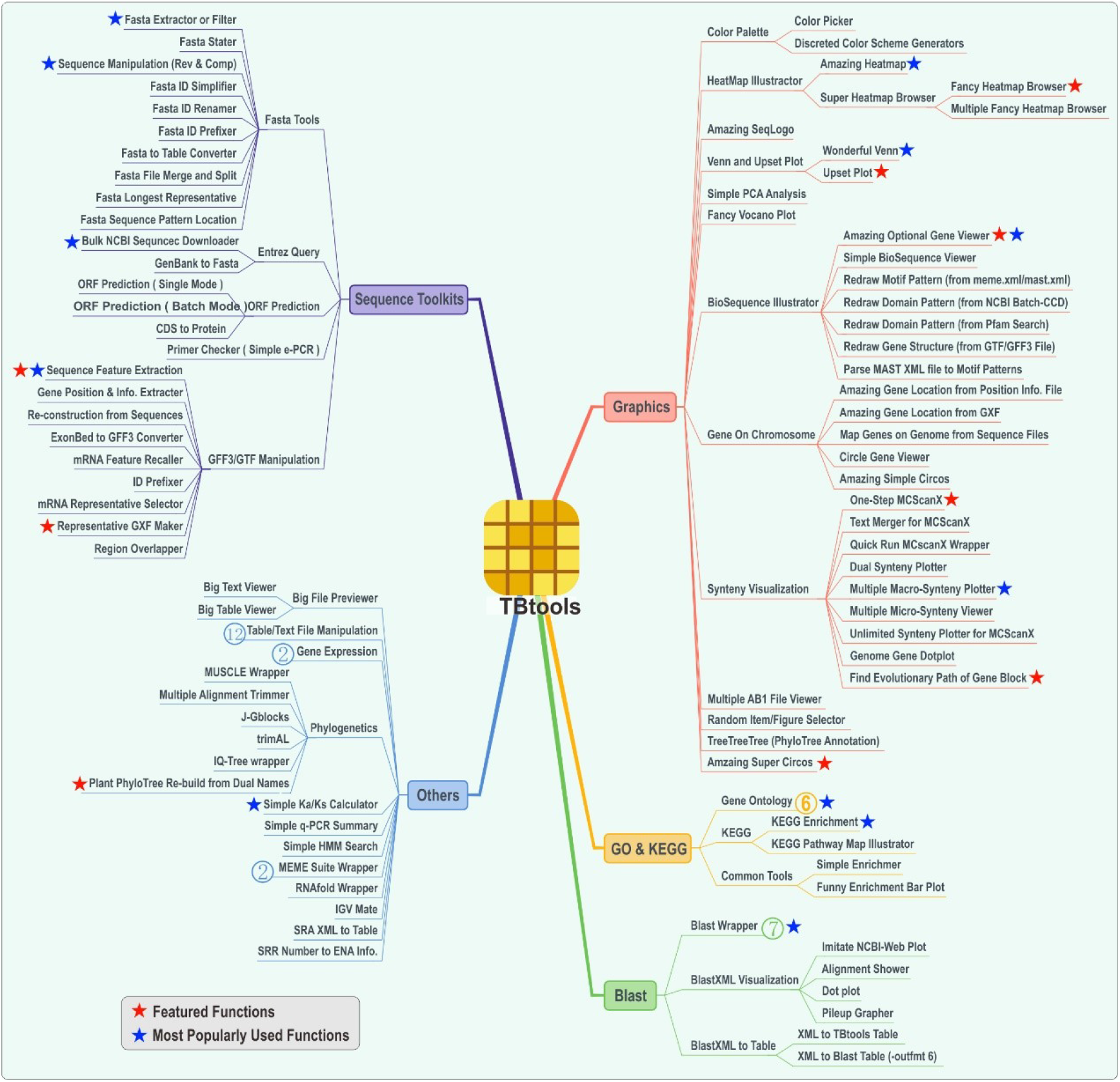
Functional outline of TBtools. There are five core catalogs that contain over 100 functions in TBtools. Featured that are most popularly used are highlighted with stars. Numbers in circles denote the number of subfunctions contained under each function.

### Developed to interactively present data

Data visualization and presentation are indispensable parts of bioinformatic analyses. In contrast to regular graph generators, which usually produce uneditable figures, TBtools generates interactive graphs full of editable features. A newly developed plotting engine named “JIGplot” (<u>J</u>ava <u>I</u>nteractive <u>G</u>raphics) is incorporated into TBtools, enabling rapid modifications to various graph features (Fig. 2). Users can easily trigger the preset functions by double-clicking on graphic elements or right-click to modify visual aspects, such as color, shape, stroke size, text, etc. Different graphic panels (e.g., phylogenetic tree, sequence alignment, gene structure, heatmaps) can be conveniently combined or arranged according to the user’s needs. Thus, the interactive nature of JIGplot allows users to simultaneously visualize data and prepare publishable graphs, without investing extra time re-plotting prestructured datasets. Another distinguishing feature of JIGplot is its unique function for coordinate transformation. For example, the coordinates of graphs can be easily switched from cartesian to polar, which allows more information to be displayed through circular plotting (Fig. 2). Besides, all graphs prepared in TBtools can be exported in both high-resolution bitmap and vector formats to allow maximum flexibility for the user’s end.

**Figure 2.**
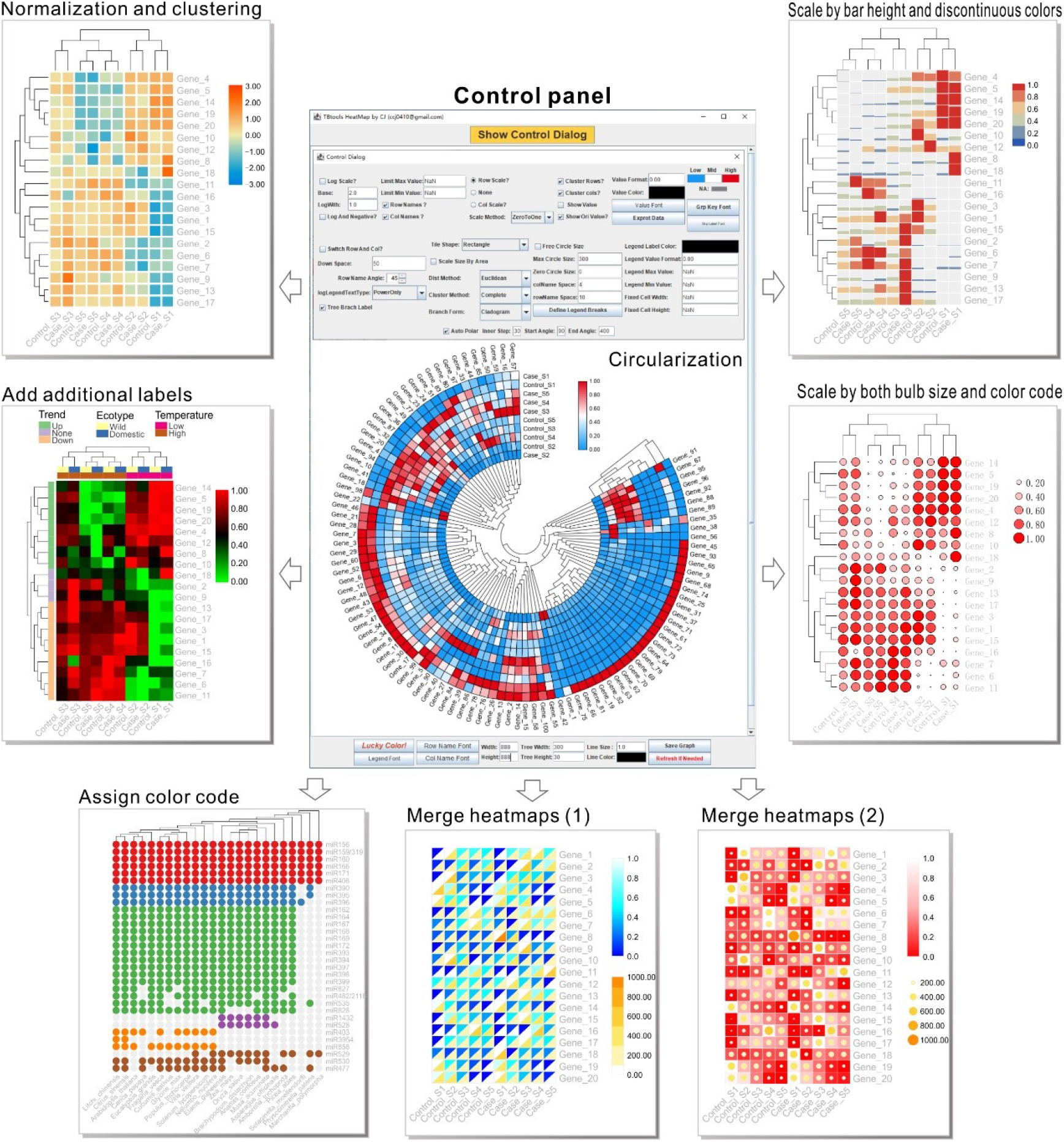
The powerful plotting engine ‘JIGplot’ displays great interactability in TBtools. A variety of heatmaps generated by TBtools are used for the demonstration of its great interactive ability and functional diversity.

## Overview of TBtools

### Function outline

All the functions in TBtools are grouped under five main catalogs (Fig. 1): (1) Sequence Toolkits. This includes all functions related to sequence management, such as sequence extraction, format conversion, and ORF prediction. (2) BLAST. A simplified BLAST wrapper was implemented in TBtools (Camacho et al., 2009); this allows sequence comparison on a local machine with optimized settings. Functions facilitating BLAST result visualization and management are also provided under this menu. (3) GO&KEGG. Enrichment analysis is a common approach to investigate the biological significance of specific gene sets. In TBtools, we have developed several functions for enrichment analysis of gene ontologies (GO) (Ashburner et al., 2000) and KEGG pathways (Kanehisa and Goto, 2000). Result files can be quickly visualized with an easy-to-use bar plot function, and sub-grouped genes to each specific GO/KEGG term can be extracted for further analyses. (4) Graphics. A wide variety of graphs can be rapidly prepared in TBtools, including Venn diagrams, heatmaps, Circos graphs (Connors et al., 2009), eFP figures (Winter et al., 2007), and so on. Most graphic features (e.g., color, shape, label) can be personalized according to individual preference. (5) Others. the Other class contains many additional functions that are commonly used during big-data exploration. It includes tools used for previewing big files, editing phylogenetic trees, and calculating Ka/Ks values.

### Stellar functions

Functions in TBtools have been improved, expanded, and optimized based on demand and feedback from our user base which includes over 15,000 stable users worldwide, many of whom are actively involved in the improvement of TBtools. A series of stellar functions are widely used by users; the results and/or graphs prepared using TBtools have been featured in hundreds of peer-reviewed publications. These functions include eFP Browser, Interactive Heatmap, Simple Circos, Gene Family Tool, and many more (Fig. 3). Although most of these functions were not originally invented in TBools, they are optimized, upgraded, and simplified according to the philosophy of “simple is best”, ensuring accessible usage for biologists.

**Figure 3.**
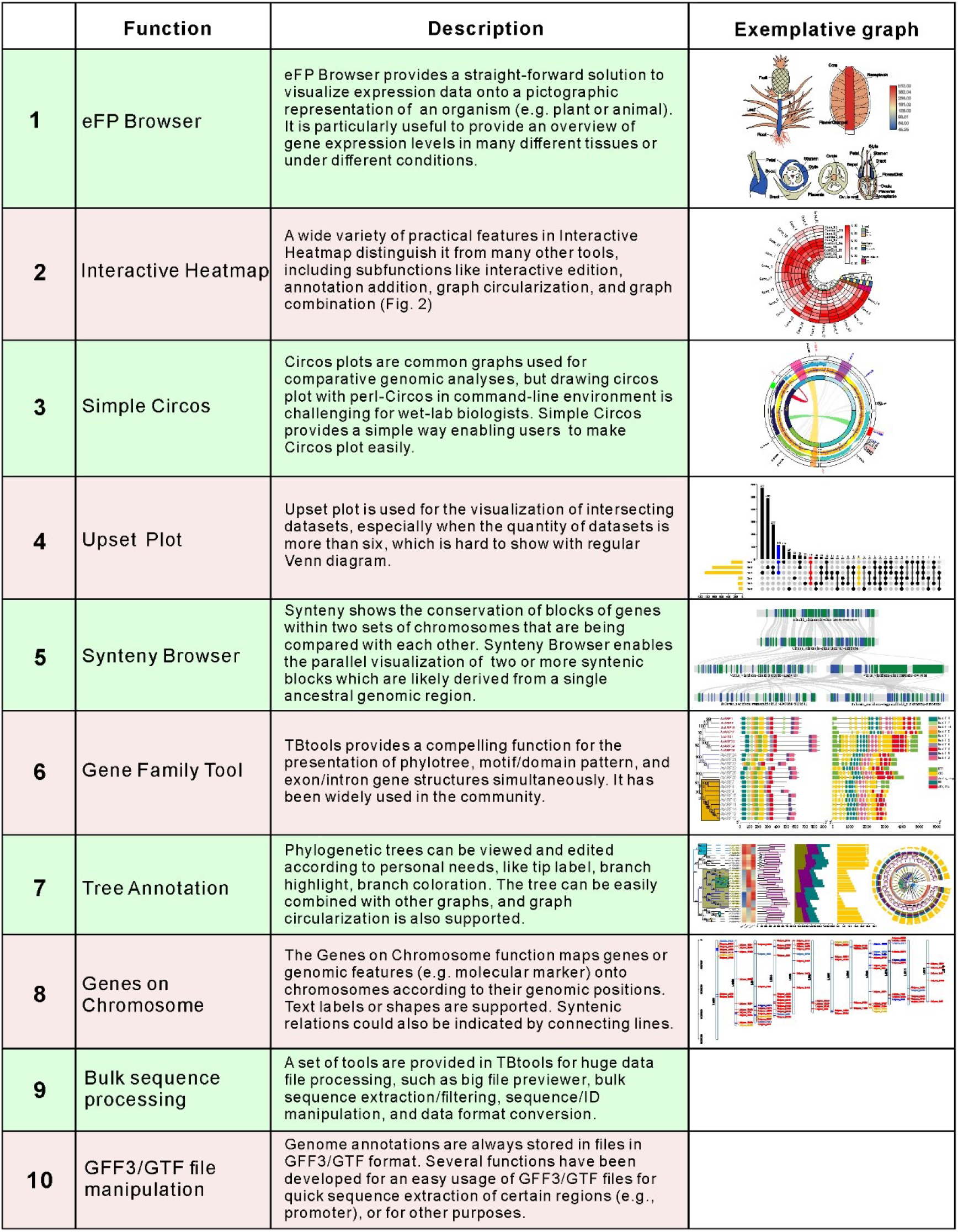
A description of prevalently used (top 10) functions in TBtools

## Conclusion

High-throughput sequencing techniques have generated vast amounts of biological data. For efficient and effective handling of this data, we have developed TBtools, a user-friendly toolkit integrated with a large number of functions with an emphasis on bulk data processing and visualization. Its robustness has been validated by tens of thousands of users, making it a handy and useful toolkit for biologists.

## Acknowledgments

This work was funded by the National Key Research and Developmental Program of China (2018YFD1000104). This work is also supported by awards to R. X., Y. H. and H. C. from the National Key Research and Developmental Program of China (2017YFD0101702, 2018YFD1000500, 2019YFD1000500), the National Science Foundation of China (#31872063) and the Special Support Program of Guangdong Province (2019TX05N193). Support to M. H. F. comes from the NSF Faculty Early Career Development Program (IOS-1942437). We thank all labmates in the Xia lab and He lab for their generous help. We are also grateful for the kind advice from 15,000+ TBtools users, especially the >30 advanced users.

